# Ontogeny shapes individual specialization

**DOI:** 10.1101/2023.04.17.537142

**Authors:** Anne G. Hertel, Jörg Albrecht, Nuria Selva, Agnieszka Sergiel, Keith A. Hobson, David M. Janz, Andreas Mulch, Jonas Kindberg, Jennifer E. Hansen, Shane C. Frank, Andreas Zedrosser, Thomas Mueller

**Affiliations:** Behavioural Ecology, Department of Biology, Ludwig-Maximilians University of Munich, Planegg-Martinsried, Germany; Senckenberg Biodiversity and Climate Research Centre (SBiK-F), Frankfurt (Main), Germany; Institute of Nature Conservation, Polish Academy of Sciences, Adama Mickiewicza 33, 31120 Krakow, Poland; Departamento de Ciencias Integradas, Facultad de Ciencias Experimentales, Centro de Estudios Avanzados en Física, Matemáticas y Computación, Universidad de Huelva, 21071 Huelva, Spain; Estación Biológica de Doñana, Consejo Superior de Investigaciones Científicas, 41020 Sevilla, Spain; Environment and Climate Change Canada, Science and Technology, Saskatoon, SK, Canada; Department of Biology and Advanced Facility for Avian Research (AFAR), University of Western Ontario, London, ON, Canada; Department of Veterinary Biomedical Sciences, Western College of Veterinary Medicine, University of Saskatchewan, 52 Campus Drive, Saskatoon, Saskatchewan S7N 5B4, Canada; Institute of Geosciences, Goethe University Frankfurt, 60438 Frankfurt (Main), Germany; Norwegian Institute for Nature Research, Trondheim, Norway; Department of Wildlife, Fish, and Environmental Studies, Swedish University of Agricultural Sciences, Umeå, Sweden; Department of Natural Sciences and Environmental Health, University of South-Eastern Norway, Bø, Norway; Institute of Wildlife Biology and Game Management, University of Natural Resources and Life Sciences, Vienna, Austria; Department of Biological Sciences, Goethe University Frankfurt, 60438 Frankfurt (Main), Germany

**Keywords:** Dietary specialization, heritability, maternal effects, social learning, trophic position, trophic niche, omnivore, stable isotopes, nitrogen-15, *Ursus arctos*

## Abstract

Individual dietary specialization, where individuals occupy a subset of a population’s wider dietary niche, is a key factor determining a species resilience against environmental change. However, the ontogeny of individual specialization, as well as associated underlying social learning, genetic, and environmental drivers, remain poorly understood. Using a multigenerational dataset of female European brown bears (*Ursus arctos*) followed since birth, we discerned the relative contributions of environmental similarity, genetic heritability, maternal effects, and offspring social learning from the mother to individual specialization. Individual specialization accounted for 43% of phenotypic variation and spanned half a trophic position, with individual diets ranging from omnivorous to carnivorous. The main determinants of dietary specialization were social learning during rearing (13%), environmental similarity (9%), maternal effects (11%), and permanent between-individual effects (8%), whereas the contribution of genetic heritability was negligible. The trophic position of offspring closely resembled the trophic position of their mothers during the first 3-4 years of independence, but this relationship ceased with increasing time since separation. Our study shows that social learning and maternal effects are as important for individual dietary specialization as environmental composition. We propose a tighter integration of social effects into future studies of range expansion and habitat selection under global change that, to date, are mostly explained by environmental drivers.

## 1. INTRODUCTION

Among individuals of the same species, ecological niche variation is common and may occur when availability of food resources or habitat structure change across the species’ range. Individual variation is key for driving species resilience in response to shifting resource availabilities in a rapidly changing world, and may ultimately determine local persistence or extinction of species ^1^. Ecological generalists, species with a wide ecological niche, also seem to exhibit more individual specialization (i.e. between-individual variation of niche) ^2^ and are likely particularly well adapted to persist under shifts in resource availability or composition, enabling them to occupy larger distributional ranges than ecological specialists ^3^. Inter- and intraspecific competition, predation and ecological opportunity alter resource availability and have been identified as the main ecological drivers explaining variation in the degree of individual dietary specialization among populations ^4^. However, how individual variation in dietary specialization emerges and is maintained within populations has, to our knowledge, not been quantified in the wild.

In the fields of behavioral and evolutionary biology, individual variation is measured as the variance attributed to permanent between-individual differences, while the sources of variation can be quantified using complex hierarchical models (e.g., “animal model”)^5^. In principle, three sources of variation are commonly considered ^5, 6^: variation in the environment ^7^, additive genetic effects from which trait heritability can be estimated ^8, 9^, and parental (especially maternal) effects ^10, 11^. In addition, individual variation can be maintained through social learning during ontogeny^12^, an aspect that, to our knowledge, has rarely been integrated into animal models (but see ^13, 14^). Here, we provide the first study to attribute individual variation in dietary niche to its sources.

Differences in the environment, in terms of habitat composition and associated availability of particular food resources, are generally considered the main cause of individual variation in dietary specialization ^15^. This is particularly true in range-resident species, where individuals occupy a subset of the population’s range and individual home ranges vary in resource availability ^16^. However, beyond the environment, resource preferences have been suggested to be genetically heritable and determined through genes inherited from both mother and father, where more closely related individuals share more similar diets than distantly related individuals ^15, 17^. Additionally, parental phenotypes may also affect offspring phenotypes in ways other than genetic heritability ^10, 11^. Maternal effects are more commonly studied because mothers often have unilateral control over offspring development ^10^, especially in mammals, however, paternal effects are plausible in species with paternal care. Maternal effects on offspring behavior have been suggested to be lifelong, they have either a genetic or environmental basis and summarize the cumulative influence of many different proximate maternal effects, include pathways such as provisioning rates, milk production, in-utero hormone transfer, and epigenetics ^10^. Statistically, maternal effects account for similarities in dietary niche among offspring of the same mother (fitted as a random intercept for mother identity) ^13, 14^, but not for the similarity of dietary niche between mother and offspring. The latter would be an example where the maternal trait affects the offspring’s trait, which statistically can be clearly differentiated from other maternal effects ^13, 14^. Similarities between the dietary phenotypes of mothers and their offspring indicate social learning of resource preference or competence to secure a resource by the offspring from the mother during early ontogeny ^18, 19, 20, 21, 22^. Social learning is therefore an additional pathway by which individual variation can be maintained. It is reasonable to assume that the effects of social learning during rearing will weaken later in life through individual experiential learning ^23^.

Attributing variation in diet to the individual level, to isolate its sources and to identify developmental drivers of diet preferences in the wild requires multigenerational datasets of repeated measures of the diet of individuals throughout their life ^5^. We used a unique 30-year longitudinal dataset of 71 female Scandinavian brown bears (*Ursus arctos*) of known mothers with repeated annual isotopic estimates of trophic position to study, for the first time, the sources of individual dietary specialization in the wild. Brown bears are ecological generalists with a distribution range spanning the northern hemisphere from tundra to deserts, paralleled by extensive variation in diet. Populations range from tracking food resource pulses, such as spawning fish ^24^, scavenging on ungulate carcasses or preying on ungulate calves ^25^, or feeding extensively on invertebrates, to populations using primarily fruiting plant-based diets ^26, 27^. Given such extreme dietary plasticity, it is not surprising that great dietary variation has been found within populations ^28, 29^; however, the determinants and ontogeny of this variation at the individual level remain largely unknown. In ecology, differences in diet are often primarily attributed to differences in resource availability and abundance. Even within populations inhabiting a continuous biome, home-range scale variation in habitat composition ^30^ can lead to variation in resource availability. Brown bears maintain non-territorial home ranges and the most parsimonious source of individual specialization is, therefore, heterogeneity in the environment. It is further plausible that individual specialization in brown bears is genetically heritable. For example, while body size is determined largely by resource availability in the environment, it has also been shown to be genetically heritable in our study population ^31^, suggesting greater similarity among closely related individuals also in other linked traits, such as trophic position. In addition, maternal effects could shape individual specialization in brown bear offspring. As a potential pathway, milk quantity or quality ^32^ can vary among females due to genetic differences and/or differences in their home range quality, leading to consistently larger or smaller offspring from the same mother, which in turn could cause similarities in trophic position among siblings. Last, brown bears live a solitary lifestyle except for the period of offspring rearing involving up to three years of maternal care ^33^, after which female offspring often settle close to their mother’s home range ^34^. In their first years of life, bear cubs accompany their mother, and so it is reasonable to predict that brown bear offspring learn their dietary niche from their mothers. If mothers differ in dietary niches, these differences may be maintained in the population through offspring social learning from the mother (hereafter “social learning”).

Individual trophic position is one metric to assess individual specialization along a continuum from a more plant-based to a more meat- or insect-based diet. Trophic position can be estimated from the ratio of stable-nitrogen isotopes (δ^15^N) in growing tissue and reflects cumulative diet intake during the period of tissue growth. Individuals with higher trophic positions are specialized on more protein-rich diets, relative to individuals with lower trophic positions which are increasingly more herbivorous. Trophic position rarely provides information on specific dietary items^2, 18^ or individual variation in niche breadth^35^ but rather quantifies the consumption of animal matter relative to other individuals in a population of omnivores. We analyzed annual trophic positions from δ^15^N values in brown bear hair keratin ^36^. Hair δ^15^N represents a dietary integration of about a month (i.e., a growing hair in June reflects the diet intake since May ^37^). Bear hair is annually renewed through molting in June, regrows over the summer and fall and stops growing during winter hibernation (**Fig 1A**, ^38, 39^). Guard hair samples collected in spring therefore reflect annual estimates of the cumulative protein intake of individuals during the previous active foraging season ^38^.

**Figure 1.**
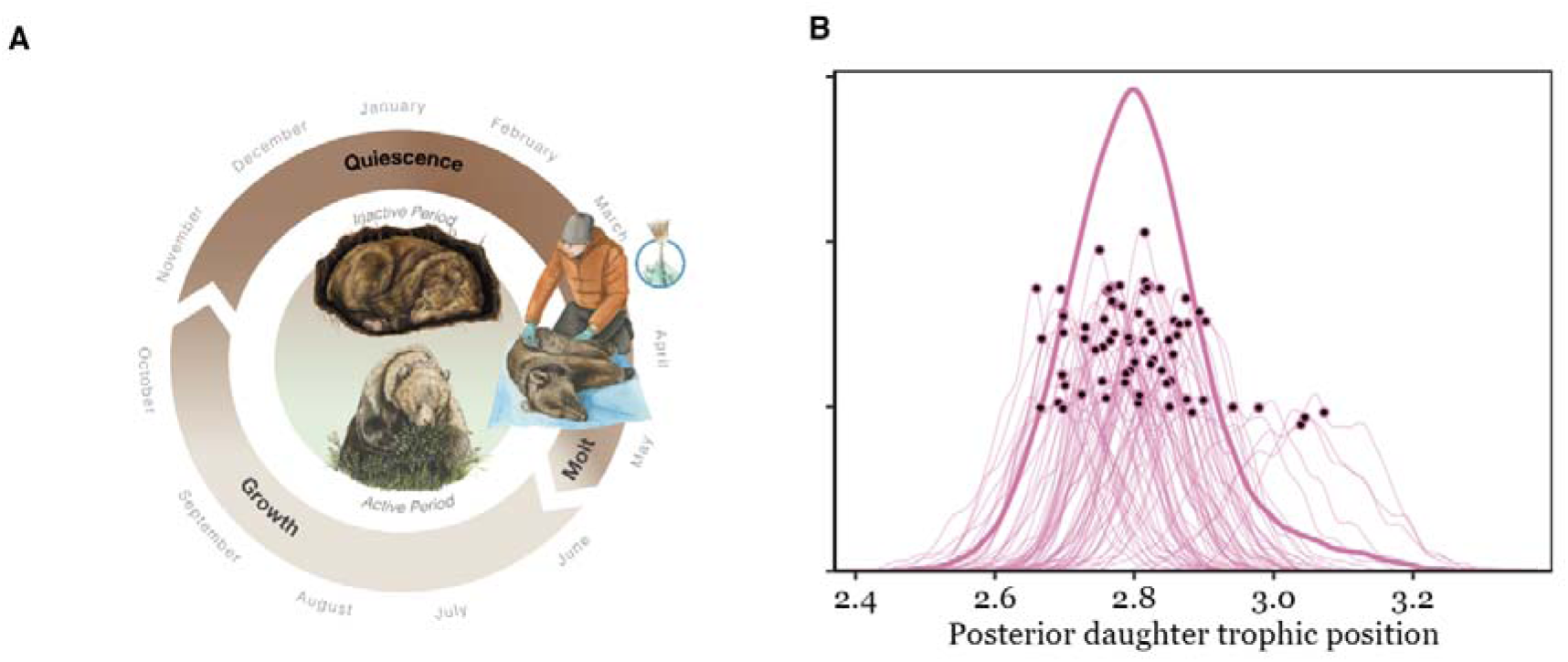
**A**) Bear hair generally grows from June until October and stable-nitrogen isotopes (δ15N) reflects cumulative diet intake during the period of hair growth. The quiescent phase, when hair ceases growing, lasts through hibernation, followed by emergence from the winter den and molting in late May-early June. Hair samples were taken during bear captures in April - June and reflect the bears’ diet in the previous year; **B**) Posterior distribution of the population trophic niche (bold line) and individual dietary niches indicated by each individual’s posterior trophic position (modelled distribution with individual posterior medians indicated by black dots). Scientific illustration by Juliana D. Spahr, SciVisuals.com.

Using repeated samples of known mother-daughter pairs, we first estimated the extent of dietary specialization as permanent individual variation and second fitted a spatially explicit Bayesian hierarchical model (i.e., ĺanimal modeĺ ^7, 40, 41^) to quantify its sources. Specifically, we accounted for environmental similarity, with pairwise habitat similarity in individual bear home ranges encompassing the proportion of mature habitat, disturbed habitat, and habitat diversity. We further accounted for genetic heritability with a pedigree, for maternal effects by incorporating the mother’s ID as a random effect, and for social learning as the fixed effect of a mother’s trophic position on her offspring’s trophic position. We allowed the effect of social learning to shift with time since offspring gained independence to account for individual learning later in life. We determined maternal trophic positions from a population-wide model accounting for sexual dimorphism, age, and permanent between-individual variation in diet (**Supplement S3**). We validated that a similarity between offspring and mother trophic position reflected social learning during rearing, and found that their trophic positions were highly correlated when together in the first year of life (**S4**). As we were interested in lifelong variation of dietary niche, and male offspring were only monitored for a short period after family breakup, we focused on individual specialization of female offspring. In the supplementary material we provide an additional analysis of the relationship between maternal and male offspring trophic position in the first four years after family breakup (**S8**) and of the relationship between paternal trophic position and offspring trophic position (**S9**). We also provide an alternative analysis accounting for spatial correlation via a spatial distance instead of an environmental similarity matrix (**S6**), as well as a reduced model excluding the effect of environmental similarity to test whether spatial and genetic effects were confounded in philopatric female bears (**S7**).We validated our effect of social learning by refitting the model to a reduced dataset with observed maternal trophic positions during rearing, instead of modeled-averaged maternal trophic positions (**S10**).

## 2. RESULTS

We analyzed annual trophic positions from 213 hair samples collected from 71 female brown bears born to 33 unique mothers (median 2 daughters; range 1-7 daughters per mother). Repeated sampling (median 3 years; range 1-11 years) showed that female trophic position was unaffected by age (median [mean, 89% equal tails credible interval] explained variance = 1% [1%, 0 – 4%]; see also **Fig S3A** and **Table S1**) and that individuals showed long-term individual specialization, accounting for 48% [47%, 31 – 61%] of the total variance in trophic position (**Fig 2**, *Basic model*). Individual variability in trophic position spanned half a trophic position ranging from 2.7 to 3.1 for individual females (**Fig 1B**), which is equivalent to the difference between an omnivore feeding on a mix of plants and animal prey and a carnivore feeding predominantly on animal prey.

**Figure 2.**
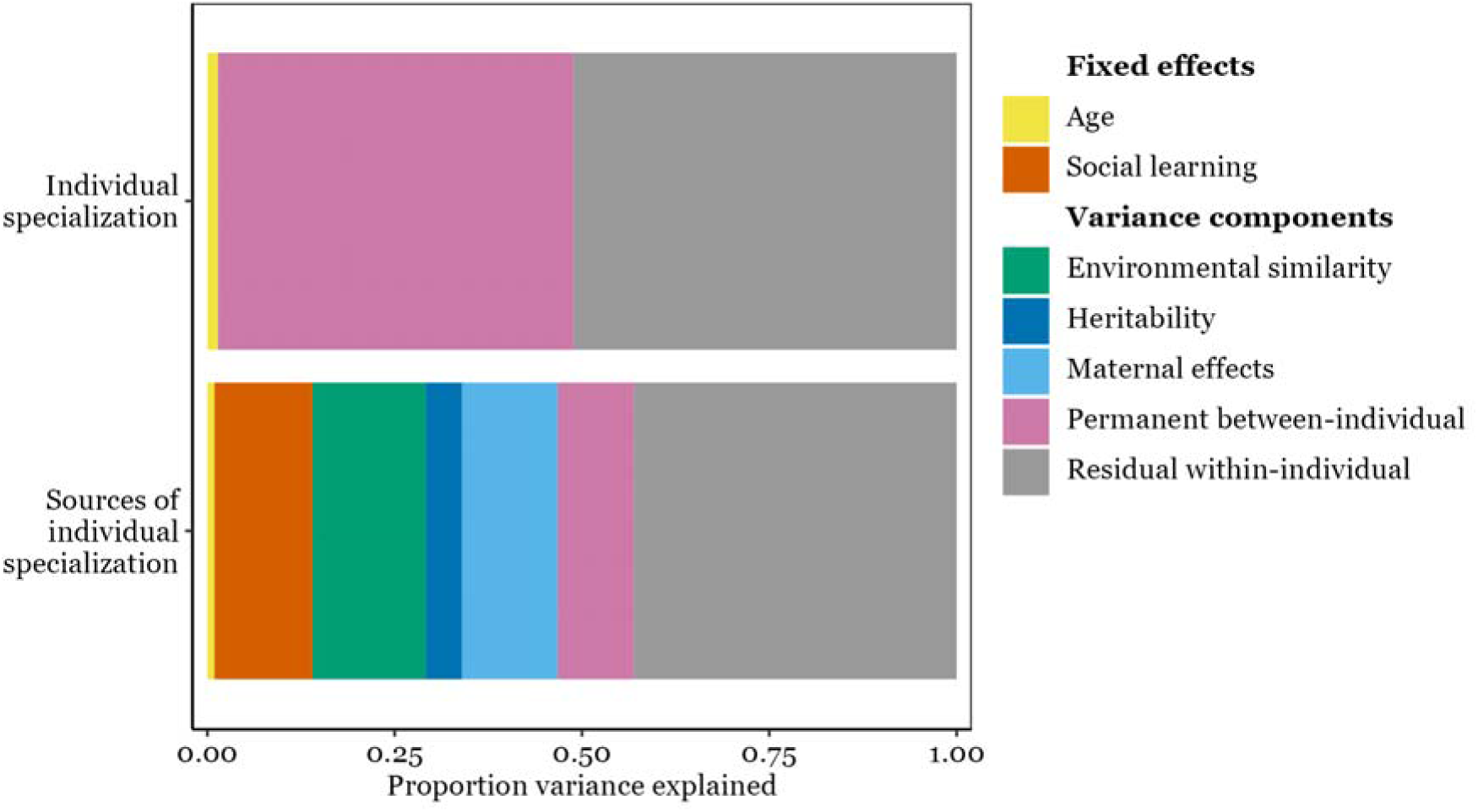
Individual specialization accounted for 48% of the phenotypic variation in trophic position of female brown bears in Central Sweden. Trophic position did not change with age. We determined the proportion of variance (mean of the posterior distribution) explained by different sources of individual specialization: Offspring social learning from the mother, environmental similarity, genetic heritability, maternal effects, permanent between-individual effects, and residual within-individual components.

Individual specialization was primarily driven by social learning, environmental similarity, and maternal effects. Maternal trophic position dynamic over time since separation accounted for 13% [13%, 5% - 23%] of the total phenotypic variation in trophic position, while environmental similarity accounted for 9% [15%, 0.1 - 49%]. Additionally, maternal effects accounted for 11% [13%, 0.5% - 30%] of variation in trophic position, indicating that siblings (full and half) of the same mother were more similar in trophic position throughout life compared to non-siblings. A remaining 8% [10%, 0.3 – 26%] of variance in trophic position was attributed to permanent between-individual effects (**Fig 2**). Genetically more closely related individuals (including paternal half-siblings, aunts, uncles or cousins) did not share a more similar trophic position (2% [5%, <0.1% – 17%] of variance explained), providing no evidence that dietary specialization could be heritable in this population (**Fig 2**, see **S11** for the full summary table).

After family breakup, female offspring initially maintained a similar trophic position to their mother (Pearson’s *r* = 0.66 in the first two years after separation), which gradually became more dissimilar over time (Pearson’s *r* = 0.31 in year 3 - 4 after separation, **Fig 3A**). In the first years, offspring of more carnivorous mothers also had a high trophic position while offspring of less carnivorous mothers had a lower trophic position. About five years after the separation from the mother, this correlation ceased to exist. Additionally, daughters of the same mother shared similar dietary niches, with siblings occupying consistently lower or higher trophic positions (**Fig 3B**). Bears inhabiting home ranges with a similar composition of mature and disturbed forest, as well as a similar habitat diversity in the home range, also had more similar trophic positions. The distance between pairwise home range centroids ranged from 0.7 to 172 km with a median pairwise distance of 48 km and individuals living in closer proximity had a more similar trophic position than individuals living farther apart (**S6**). However, spatial distance and maternal effects seemed to be confounded in this female philopatric species (**S7**): after excluding spatial distance, social learning and maternal effects, but not heritability, explained more variance in trophic position, indicating that spatial proximity and maternal effects are confounded because settlement home ranges of philopatric daughters are often close in space to their mothers forming so called matrilineal assemblages. In a separate analysis (**S8**) we also showed that the relationship between maternal and offspring trophic position in the first years after family breakup was not sex-specific. Both male (n = 31, Pearson correlation coefficient = 0.4) and female (n = 69, Pearson correlation coefficient = 0.45) offspring trophic positions resembled their mother’s trophic position in the first 4 years of independence, corroborating our findings that initial social learning determines foraging behavior in the early years after family breakup. Conversely, paternal trophic position had no effect on offspring trophic position in the first four years of independence (Pearson correlation coefficient = 0.13, **S9**). While the modelled maternal trophic position correlated strongly with the observed trophic position in any given year (**S3, Fig S3B**), social learning explained even more of the phenotypic variance in daughter trophic position (22% [8% - 27%] instead of 13 %, **S10**) when fitting the observed maternal trophic position during rearing, instead of the modelled posterior average maternal trophic position to a reduced dataset (62 hair samples collected from 38 daughters). Our estimates of offspring social learning from the mother are therefore likely conservative and may underestimate the true effect of social learning on individual specialization.

**Figure 3.**
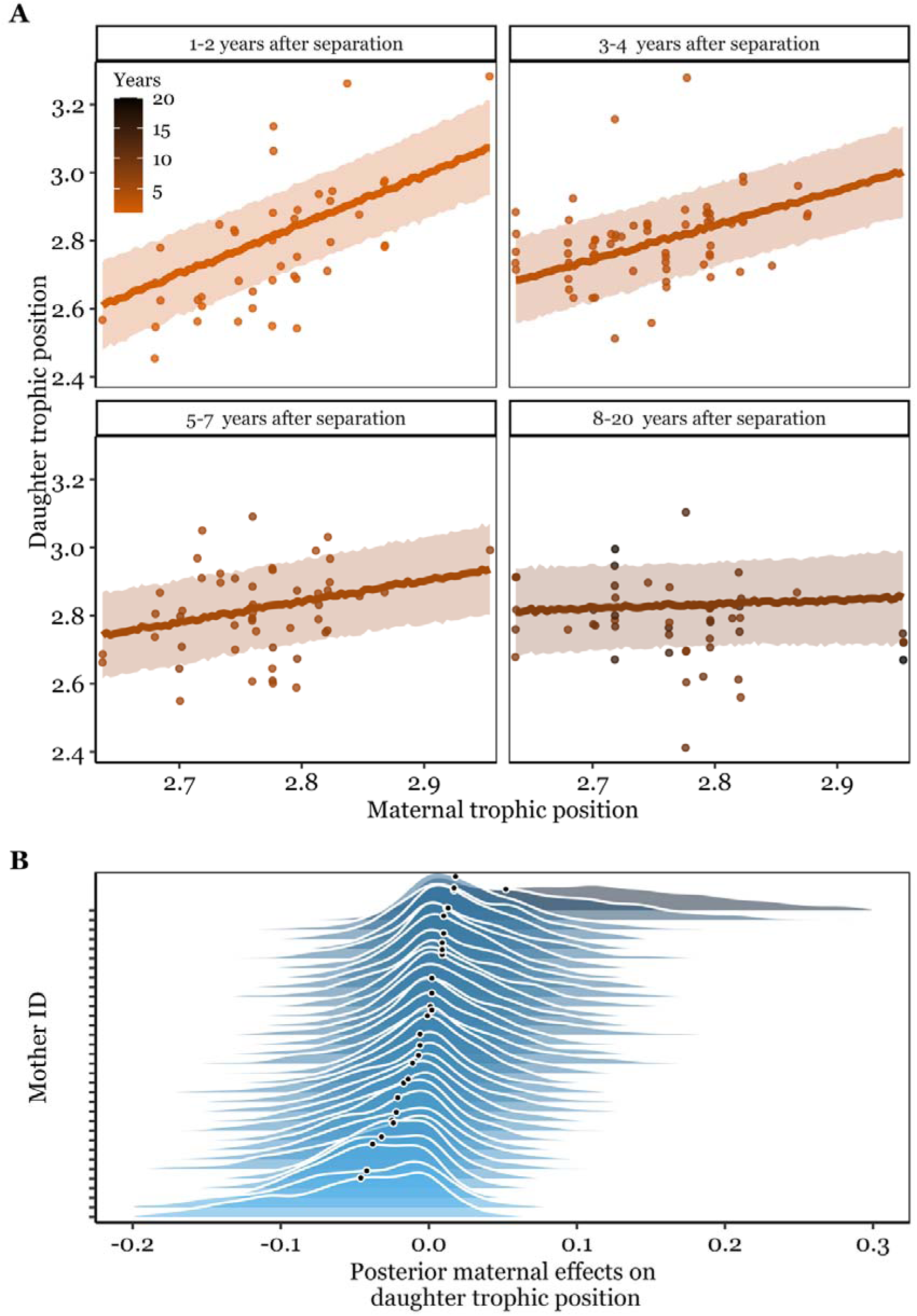
**A)** Relationship between female brown bear trophic position and their mother’s trophic position over number of years since separation, which occurs at 1.5 – 2.5 years of age in our population. The daughter’s trophic position resembled their mothers’ trophic position in the first years after separation but this similarity ceased after 4 years. Lines indicate predicted posterior mean estimates with ribbons corresponding to the estimated standard error, raw data are shown as points. **B)** Additional maternal effects (e.g., maternal genotype or maternal environment) explained further similarities in trophic position among daughters of the same mother (and differences between daughters of different mothers).

## 3. DISCUSSION

Our multigenerational dataset reveals unique insights into the ontogeny of individual dietary specialization along a continuum from a more herbivorous to a more carnivorous diet in a long-lived omnivore. Specifically, the foraging strategy of offspring was intimately tied to the foraging strategy of their mother, a relationship that lasted up to four years after independence. We interpret this relationship as evidence that social learning plays an important role in shaping an individual’s dietary specialization. Five years into independence, the similarity between the trophic position of mothers and daughters slowly faded, likely due to individual learning and experience during solitary life. In addition, offspring of the same mother also shared similarities in their trophic position, potentially mediated through maternal genetic or environmental effects on body size ^31^. Additive genetic effects on the other hand were insignificant providing no evidence for heritability of dietary specialization in this population. Similar to the effect of social learning from the mother fading over time, additive genetic and maternal effects could be life-stage specific with maternal effects being more influential in juveniles^13^, although evidence for this is mixed ^10^. We were not able to quantify life-stage specific heritability and maternal effects due to sample size limitations. In general, previous ecological studies have mainly concentrated on resource availability as the main driver of resource selection ^42^ and individual specialization ^4^. However, our results show that, within populations, the environment is only one of several components shaping individual variation in dietary niche. We conclude that social learning of maternal dietary preferences during early-life ontogeny and maternal effects (i.e., maternal genotype and environment), which together explained about 24% of the variation in trophic position, play a pivotal role in spreading and maintaining feeding strategies within populations, even in species with otherwise solitary lifestyles. In addition, variation linked to between-individual effects (in our study 8 %) could be associated to either uncontrolled variation in resource availability in the environment (i.e., ecological opportunity ^4, 35^) or individual differences in resource preference. The latter could, for example, be caused by differential individual learning and demonstrates the potential for behavioral innovation in this population. Ultimately, between-individual variation in dietary specialization allows populations to adapt to changes in resource availability, such as, new invasive prey or declines in food items due to climate change.

Our findings are particularly relevant for species in which dietary specialization impacts individual fitness ^20, 35, 43^. For example, protein-rich diets may promote greater offspring survival or mass gain ^44^. Social learning in general, therefore, presents an important, yet understudied, pathway by which alternative behavioral strategies can establish and spread more rapidly within populations than by genetic evolution alone ^45^. Species more adept in social learning of dietary strategies may therefore show greater behavioral variability at the population level, which could give them an advantage when adapting to changing environments due to landscape modification or urbanization, climatic variations or global change in general. Moreover, there is evidence that the strength of social learning in shaping individual phenotypes is not only species-specific, but can also vary among populations or individuals of the same species ^12, 46^.

Our research also points to several aspects of social learning that warrant future research. First, there is little information on whether maternal care and social learning tend to be more prevalent in species or populations with greater dietary specialization. There is some evidence that within populations, dietary generalists (i.e. those with a wider dietary niche) seem to provide more intense parental care ^47^ than their conspecific dietary specialists (i.e. ones with a narrower dietary niche), but the link to social learning of foraging preferences remains unclear. Second, while generalist species with a wide ecological niche have been frequently shown to be more successful under changing environmental conditions, such as urban environments or fragmented landscapes, than specialist species ^48, 49, 50^, it is currently unknown whether this success could be partially mediated by social learning. Finally, social learning could alternatively limit behavioral innovation and adaptation due to adherence to social traditions ^51^. We therefore suggest that alternative hypotheses should be evaluated that consider how social learning impacts individual specialization and in turn the adaptability of species under global change.

Our findings that dietary specialization can be socially learned and transmitted are particularly relevant for species where individual specialization is related to human-wildlife conflict ^52^. For example, the removal of single individuals which are known to cause conflict is an effective strategy to halt the spread of problematic behavior and mitigate the conflict, while minimizing the impact on species conservation goals ^52^. Foraging behavior that causes conflict with humans has also been shown to change in ursids over their lifetimes, remarking the crucial role of individuality and plasticity in behavior ^53^. Social learning from the mother of behavior ^54^, including individual specialization and foraging on anthropogenic food resources, has been previously observed in ursids ^55, 56, 57, 58^. However, none of these studies tracked offspring diet over their lifetimes or were able to simultaneously account for the mother’s diet, genetics, the environment, and other maternal effects that could explain similar patterns of individual specialization. While some of the aforementioned studies suggest either the environment or social learning as primary drivers of individual specialization, we suggest using caution in assigning causality in dietary specialization, when potentially confounding alternative sources cannot be accounted for. Specifically, in female-biased philopatric species, spatial proximity does not only encode for spatial variation in resource abundance but is also conflated with heritability and with maternal effects. In brown bears, some daughters settle close to their mother’s home range ^34^ creating spatial clusters of closely related females, so called matrilinear assemblages ^59^. Due to spatial dependence of these assemblages, it can therefore be difficult to untangle social learning from the mother from other maternal effects (i.e., maternal genotype or maternal environment) or the ambient environment. Our study population spanned over 170 km with spatial proximity explaining 50% of the total phenotypic variation in trophic position of female bears: individuals further apart tended to have more different diets. However, when replacing spatial proximity with environmental similarity among home ranges, the explanatory power was attributed to social learning and maternal effects along with the environment. Our results therefore demonstrate that individual dietary specialization is not caused by a single driver in isolation but the product of many factors, namely social learning, maternal effects, and the environment.

Our finding that social learning has a similar impact on resource selection as the environment provides important insights for a range of studies on habitat selection, dispersal, and range expansion. For example, a popular theory known as “natal habitat preference induction” suggests that dispersing animals select areas for settlement that resemble their natal habitat, even at fine habitat scales ^30^. Our results challenge the notion that habitat similarity alone drives natal settlement strategies and rather suggest that socially learned diet preferences, and hence the selection for food resources themselves, could play an important role in producing similar patterns of settlement selection like induced natal habitat preferences. Recent studies of migration and short stopover behavior in whooping cranes (*Grus americana*) have also observed that social learning rather than environmental conditions ^60^ or genetic heritability ^61^ led to the emergence and establishment of alternative migratory behavior. Similar to what our study shows with respect to individual specialization, social learning of migration strategies primarily determined behavior in early life whereas individual-experiential learning shaped behavior later in life ^62^.

## Conclusion

Drivers of individual dietary specialization are well documented among populations of the same species. However, systematic studies delineating the sources of individual specialization within populations are lacking, likely because suitable datasets including multigenerational, genetic, environmental, and life-history information are rare. We show that, in addition to the environment, social learning and maternal effects can be important sources of dietary specialization.

## 4. METHODS

### Bear hair sample collection

We collected brown bear hair samples in south-central Sweden (∼N61°, E15°) as part of a long-term, individual-based monitoring project (Scandinavian Brown Bear Research Project; www.bearproject.info). Hair samples were collected from known individuals and their offspring during captures in spring (April - June) 1993 – 2016. Bears were immobilized from a helicopter (Arnemo & Fahlman, 2011). A vestigial premolar tooth was collected from all bears not captured as a yearling to estimate age based on the cementum annuli in the root ^63^. Tissue samples (stored in 95% alcohol) were taken for DNA extraction to assign parentage and construct a genetic pedigree ^59^. Guard hairs with follicles were plucked with pliers from a standardized spot between the shoulder blades and archived at the Swedish National Veterinary Institute. All animal captures and handling were performed in accordance with relevant guidelines and regulations and were approved by the Swedish authorities and ethical committee (Uppsala Djurförsöksetiska Nämnd: C40/3, C212/9, C47/9, C210/10, C7/12, C268/12, C18/ 15. Statens Veterinärmediciniska Anstalt, Jordbruksverket, Naturvårdsverket: Dnr 35-846/03, Dnr 412-7093-08 NV, Dnr 412-7327-09 Nv, Dnr 31-11102/12, NV-01758-14). We used data of offspring after separation from their mother (main text) and of adult male and female bears (**Supplementary analysis S3**). Bear cubs are born in January or February during winter hibernation and are typically first captured together with their mother as yearlings at the age of ∼ 15 months. Cubs in this population separate from their mother during the mating season in May or June after 1.5 or 2.5 years ^64^. A hair sample taken in spring reflects the summer-fall diet of the bear in the previous active season (**Fig 1A**). Only hair samples of solitary, independent offspring taken in spring at least 10 months after separation from the mother were included in this study.

### Moose sample collection

We collected samples of the natural foods most important for brown bear in the study area (**Fig S1**), including 21 samples of moose hair (*Alces alces*), the most common meat source in the diet of brown bears in our study area ^65^ in the spring-autumn field season of 2014.

Samples were placed in a paper envelope and dried at ambient temperature.

### Stable isotope analyses

Hair samples were rinsed with a 2:1 mixture of chloroform:methanol or washed with pure methanol to remove surface oils ^66^. Dried samples were ground with a ball grinder (Retsch model MM-301, Haan, Germany). We weighed 1 mg of ground hair into pre-combusted tin capsules and combusted at 1030°C in a Carlo Erba NA1500 elemental analyser. N_2_ and CO_2_ were separated chromatographically and introduced to an Elementar Isoprime isotope ratio mass spectrometer (Langenselbold, Germany). Two reference materials were used to normalize the results to VPDB and AIR for δ^13^C and δ^15^N measurement, respectively: BWB III keratin (δ^13^C =- 20.18‰, δ^15^N = 14.31‰, respectively) and PRC gel (δ^13^C =-13.64‰, δ^15^N = 5.07‰, respectively). Measurement precisions as determined from both reference and sample duplicate analyses were ± 0.1‰ for both δ^13^C and δ^15^N.

### Bear trophic position

We calculated the trophic position of each bear hair sample relative to the average δ^15^N value of moose hair representing trophic level 2 (mean ± sd = 1.8 ± 1.26 ‰, n = 21, **Fig S1**). Bears consume most of a moose carcass, including meat, skin, and hair. Soft tissue samples of moose carcasses could not be obtained but according to the literature the ratio of δ^15^N in ungulate hair to meat ranges between 0.77‰ – 1.0‰. (see S3.1 in ^67^). We consider trophic positions calculated from moose hair representative and a correction of the δ^15^N moose hair signature would only add an arithmetic correction but not change the distribution of bear trophic positions. Trophic position is calculated as the discrepancy of δ^15^N in a secondary consumer and its food source divided by the enrichment of δ^15^N per trophic level, plus lambda, the trophic position of the food source (e.g. 1 for primary producers, 2 for primary consumers, 3 for secondary consumer, 4 for tertiary consumers) ^68^. We used an average trophic enrichment factor of 3.4‰ ^68^ and added a lambda of 2 given the moose baseline trophic position as a strict herbivore.

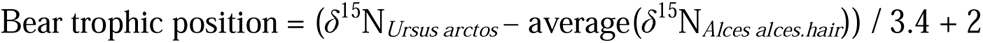

Under an omnivorous diet including the consumption of herbivores (in particular moose but also ants such as *Formica* spp*., Camponotus herculeanus* with average δ^15^N indistinguishable from moose, **Fig S1**), bear trophic position values were expected to fall between 2 and 3. Values approaching 4 indicate a trophic enrichment through consumption of other omnivorous or carnivorous animals.

### Sources of individual variation in trophic position

#### Environmental similarity

Resources may not be distributed evenly in space. For moose, population density and hunting quotas (which determine availability of slaughter remains) vary across the study area. For ants, the availability of old forests and clearcuts determine their abundance ^69^. Furthermore, brown bear daughters are often philopatric with limited dispersal and settle close to their mother’s home range ^34^. The median dispersal distance of daughters, namely the distance between natal and settlement home range centroids in this study was 8.56 km (range 1.4 – 28.8 km). Genetic, spatial, and social learning effects may therefore be confounded with related bears occupying adjacent ranges with similar resource availability. Elsewhere, accounting for environmental similarity through spatial autocorrelation in animal models has revealed that a major portion of variance may be attributed to environmental similarity rather than genetic heritability ^7, 40, 70^, but see also ^71^. Here, we accounted for environmental similarity by extracting habitat composition in each bear’s lifetime home range (n = 71). We constructed home ranges using a 95% kernel density estimator (kernelUD function) in the R package adehabitatHR ^72^ using the reference bandwidth as smoothing parameter (option “href”). Bears were monitored for a minimum of 1000 GPS locations (n = 47) or were located via VHF on at least 25 days (n = 24). Median lifetime home range size was 256 km^2^, which is comparable to a circle with an 18 km diameter. We used a Corine landcover map (25 m resolution) which we updated annually with polygons of newly emerged clearcuts (data obtained from the Swedish Forest Agency). We extracted home range habitat composition in the year when diet was assessed. When individuals were monitored for multiple years, we extracted the home range composition for the median year. Annual changes in home range habitat composition were negligible (**Fig S5**). We calculated the proportion of mid-aged and old forest and proportion of disturbed forest (clearcuts and regenerating young forest) within the 95% utilization distribution. Additionally, we calculated habitat diversity using the Simpson diversity index from the R package landscapemetrics ^73^. Following Thomson et al. ^40^ we calculated the Euclidean distance between scaled and centered habitat composition and habitat diversity in multivariate space, assuming equal importance of each component. Pairwise distances were scaled between 0 and 1, where increasing values indicated more similar habitat composition. In the supplementary material we provide an alternative analysis accounting for spatial autocorrelation of individual dietary niches with a pairwise spatial distance matrix (S matrix; **Supplementary analysis 6**, **Fig S6**). Spatial autocorrelation of home range habitat composition seized after 10 – 15 km (**Fig S6**).

#### Genetic pedigree

A genetic pedigree based on 16 microsatellite loci was available for the population including 1614 individual genotypes, spanning six generation ^59^. All female offspring and mothers in this study were genotyped and included in the population’s genetic pedigree. We used Cervus 3.0 ^74^ for assignment of fathers and COLONY ^75^ for creating putative unknown mother or father genotypes and sibship reconstruction (see for details ^59^).

#### Maternal identity

The study was based on a population of marked females and their offspring. Therefore, all mothers included in this study were known from observations of mother-offspring associations.

#### Maternal trophic position

Based on repeated hair samples of 115 female (n_female_ = 335) and 98 male (n_male_ = 219) bears, we fitted a linear mixed effects model for female and male bears respectively, to estimate sex-specific among individual variation in trophic position (**Supplementary analysis 3**). We modelled trophic position as a function of a quadratic relationship with age and we controlled for individual random intercepts. We concentrate on the relationship between age and trophic position as age is not confounded with among-individual effects, however we also perused the relationship between mass and trophic position in the **Supplementary analysis 2, Fig S2.** Female trophic position did not vary with age but was highly repeatable over multiple years. For all daughters, we extracted their mother’s (and father’s) trophic position as the median of the posterior distribution of their respective random intercept. The modelled posterior trophic position and the observed trophic position in a given sampling year were strongly positively correlated (Pearson correlation coefficient r = 0.78, t = 22.63, df = 336, p < 0.001).

### Statistical analysis

We applied a two-step modelling approach. First, we fitted a *basic* linear mixed-effects model to estimate individual dietary specialization as permanent between-individual variation (*V*_I_) in trophic position. For this we used repeated measures of the same individual and fitted individual random intercepts. We accounted for a nonlinear effect of age (second-order polynomial, scaled by the standard deviation). We extracted the variance in fitted values (*V*_Age_; variance explained by age), permanent between-individual (*V*_I_), and residual within-individual variance (*V*_R_) and estimated each component’s proportional contribution to the total phenotypic variance (*V*_P_ = *V*_Age_ + *V*_I_ +*V*_R_) through variance standardization ^76^.

Second, we used a spatially explicit Bayesian hierarchical model (i.e. ‘animal model’) ^5, 40, 41^ to partition permanent between-individual variance (*V*_I_) in trophic position into its sources; the fixed effects age (*V*_Age_) and social learning (*V*_SL_), and the variance components permanent between-individual variance (*V*_I_), environmental similarity (*V*_E_), additive genetic variance (*V*_A_), maternal effects (*V*_M_), and residual within-individual variance (*V*_R_). We followed the “hybrid” strategy suggested by McAdam, Garant ^14^ and tested for social learning of trophic position from the mother (“social learning”) by incorporating maternal trophic position as a fixed effect into the model. Thus, *V_M_* pools the remaining phenotypic variation of offspring trophic position that cannot be explained by maternal trophic position. We accounted for a nonlinear effect of time since separation of mother and daughter (scaled by the standard deviation), and for an interaction between maternal trophic position and time since separation to account for a decrease of the social learning effect over time. Age and time since separation were perfectly correlated: Pearson correlation coefficient > 0.99.

We calculated the total variance explained by the model (*V*_P_ = *V*_Age_ + *V*_SL_ + *V*_I_ + *V*_E_ + *V*_A_ + *V*_M_ +*V*_R_) and calculated the proportion of the total variance explained by each model component. For the fixed effects, we partitioned the variance explained into social learning over time (i.e. maternal trophic position and its interaction with time since separation) and age (i.e. the main effect of time since separation), respectively, by calculating the independent contribution of each component to the total variance explained by the fixed effects, following the approach by Stoffel, Nakagawa ^77^ adapted to a Bayesian framework (see code under ^78^). For all parameters, we report the median and mean as measures of centrality and 89% credible intervals, calculated as equal tail intervals, as measure of uncertainty ^79, 80^. We deemed explained variance proportions as inconclusive when the lower credible interval limit was < 0.001 (i.e., < 0.1%) ^81^.

All models were fitted using the R package “brms” ^82^ based on the Bayesian software Stan ^83, 84^. We ran four chains to evaluate convergence which were run for 6,000 iterations, with a warmup of 3,000 iterations and a thinning interval of 10. All estimated model coefficients and credible intervals were therefore based on 1200 posterior samples and had satisfactory convergence diagnostics with 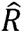 < 1.01, and effective sample sizes > 400 ^85^. Posterior predictive checks recreated the underlying Gaussian distribution of trophic position well. All statistical analyses were performed in R 4.0.0 ^86^. Primary data and code to reproduce all analyses are provided under (https://doi.org/10.17605/OSF.IO/68B9U, ^78^).

## ACKNOWLEDGEMENTS

AGH has received funding from the European Union’s Horizon 2020 Research and Innovation Programme under the Marie Skłodowska-Curie Grant agreement No 793077 and from the German Science Foundation (HE 8857/1-1). The study was further funded by the Norway Grants under the Polish-Norwegian Research Programme administered by the National Research Centre for Research and Development in Poland and the Norwegian Research Council (JA, NS, AS, and AZ; GLOBE No POL-NOR/198352/85/2013). Isotope analyses were funded through a Robert Bosch Foundation grant to TM and the GLOBE project and conducted by KAH (with assistance from Blanca X. Mora Alvarez and Geoff Koehler), DJ and AS. We thank the Scandinavian Brown Bear Research Project (SBBRP) for providing access to the data. The SBBRP was funded by the Norwegian Environment Agency, the Swedish Environmental Protection Agency, the Austrian Science Fund, and the Norwegian Research Council.

## AUTHOR’S CONTRIUTIONS

AH, JA, and TM developed the work. AZ, JK, KAH, NS and AS provided the data. AZ managed the sample collection. AS managed the hair samples database and prepared samples for stable isotope analyses by KAH. DJ provided laboratory space and resources and supervised preparatory procedures. SF constructed and provided the genetic pedigree. JH provided home range centroids. AM and JA advised to the analysis and interpretation of stable isotope data. TM, NS and AZ secure project funding. AH performed the statistical analyses with input from JA. AH wrote the manuscript with help from TM, JA, and input from all authors.

