## Supplementary material for "Ontogeny shapes individual specialization"

**Online Supplementary material:**

1. **Stable isotopic ratios in food sources**
2. **Relationship between age, mass, and trophic position**
3. **Estimate maternal and paternal trophic position**
4. **Maternal learning during rearing**
5. **Distance between home range centroids of mothers and daughters over time**
6. **Effect of environmental similarity versus spatial proximity on trophic position**
7. **Collinearity of genetic effects and spatial proximity**
8. **Effect of maternal trophic position on male and female offspring**
9. **Effect of paternal trophic position on offspring trophic position**
10. **Alternative analysis of the effect of maternal learning**
11. **Summary table of full model**
12. **Statistical power analysis**

### 1. Stable isotopic ratios in food sources

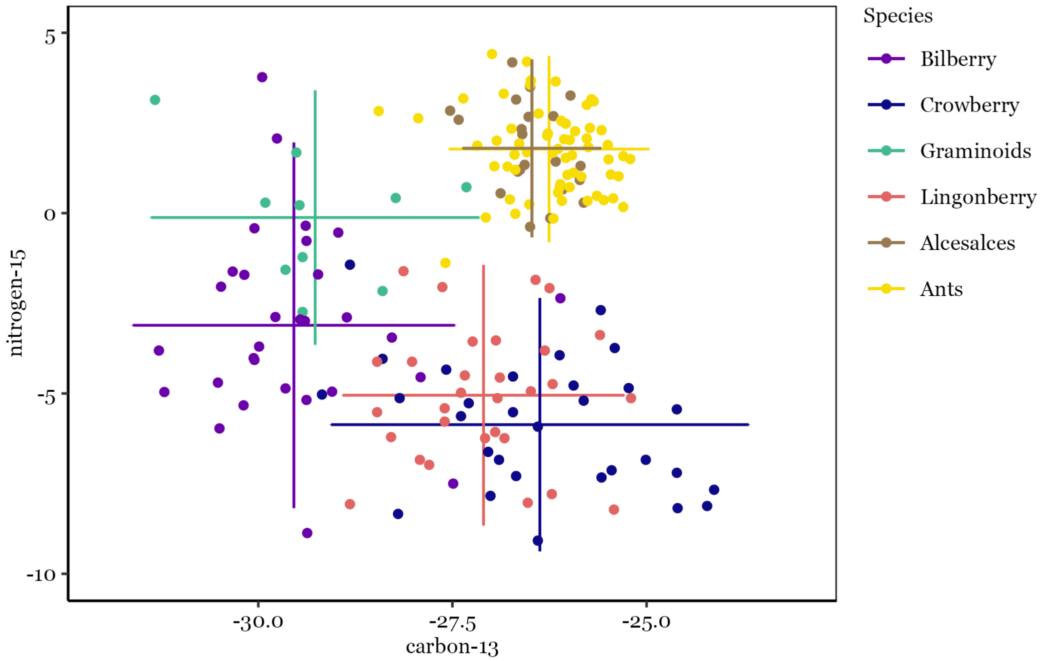


**Figure S1.** We measured stable isotopes $\delta^{13}C$ and $\delta^{15}N$ (in ‰) in the main dietary food resources of Scandinavian brown bears during the active season: bilberry (*Vaccinium myrtillus*), lingonberry (*Vaccinium vitis-idaea*), crowberry (*Empetrum sp.*), moose (*Alces alces*), and ants (*Formica spp,* and *Camponotus herculeanus*). Error bars indicate the mean ± standard deviation for each dietary item. High $\delta^{15}N$ values are characteristic of primary herbivores with moose and ant samples, indistinguishable. We used the average $\delta^{15}N$ signature of moose as a reference to calculate individual trophic positions of brown bears.

### 2. Relationship between age, mass, and trophic position

Body mass and age are both covariates which might be biologically related to trophic position. Heavier bears may be physically more capable to capture prey, however since the main prey in our study area is moose calves in their first month of life we expect that even young bears can capture prey successfully (Swenson *et al.* 2007). Further, heavier bears may be able to monopolize slaughter remains, gut piles or human killed cadavers more effectively than smaller bears. On the other hand, older bears may be more experienced and skilled in capturing moose calves and therefore have a higher predation success.

Female Scandinavian brown bears increase in mass until approximately age 10 (**Fig S2 left**). After that, spring weights of females are stable around 80 - 100 kg. There is higher variation in mass for adult male bears (150 – 250 kg). Mass and age are inherently correlated (**Fig S2 left**) and it is therefore not possible to delineate which of the two is shaping trophic position.

In the main text we are concentrating on a functional relationship between age and trophic position because the covariate mass also captures among-individual effects (bigger bears are probably bigger throughout life) while age estimates are not confounded with among-individual effects. We here show that the effect of mass on trophic position is similar for females and males, at least up to 100kg (**Fig S2 right**). For both females and males, the trophic position increases with increasing mass. However, trophic position does not increase for heavier females (>100 kg) while heavier males reach much higher trophic positions. We explain the exceptionally high trophic position values in large males by adult males killing (and consuming) brown bear cubs during the mating season to trigger estrus in females bears (Steyaert *et al.* 2012).

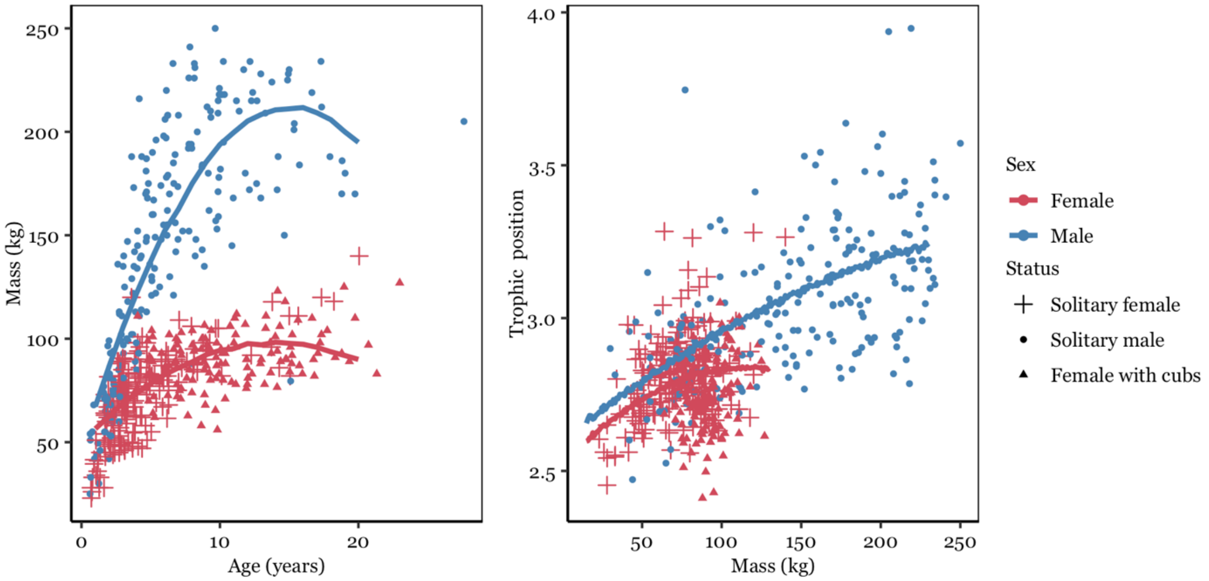


**Figure S2.** Relationship between age and mass (left) and mass and trophic position (right) in male (n = 97) and female (n = 115) Scandinavian brown bears.

#### 3. Estimate maternal and paternal trophic position

We obtained 554 hair samples of 213 individual bears (115 females and 98 males), many of which were sampled repeatedly over multiple years ($n_{female}$ = 335, $n_{male}$ = 219). We fitted two linear mixed-effects models for female and male bears respectively, to estimate sex-specific among individual variation in trophic position. We modelled trophic position as a function of a quadratic relationship with age. To estimate among-individual variance in trophic position for both male and female bears we controlled for individual random intercepts (number of repeated measures per individual for females: 1 – 11, median = 3; males: 1 – 8, median = 2). We estimated repeatability i.e. variance standardized individual variation, as among-individual variance divided by total phenotypic variance. We extracted the variance in fitted values (variance explained by fixed effects), among-individual, and residual variance and calculated Nakagawa's marginal and conditional $R^2$ (Nakagawa & Schielzeth 2013).

Annual trophic positions ranged from 2.41 – 3.95 (range $\delta^{15}N$: 3.2 – 8.42‰) indicating a wide dietary niche. On the population level, males were more carnivorous than females (**Table S1**) and with increasing age, male bear diets became more carnivorous, as indicated by a higher trophic position (**Table S1**, **Fig S3A**). Females on the other hand did not become more carnivorous with increasing age (**Fig S3A**). For an adult female at the age of 10 (**Fig S3A**) the estimated trophic position was 2.8 [2.6, 3] while for and adult male of the same age the estimated trophic position was 3.18 [2.8, 3.5]. Age accounted for 22% ($R^2marginal_{male}$ = 0.26 [0.15, 0.37]) of the variation in trophic position in male bears while it accounted for no variance of the trophic position in female bears ($R^2marginal_{female}$ = 0.01 [0, 0.02]). Therefore, for females, all explained variance in trophic position was attributed to among individual differences ($R^2conditional_{female}$ = 0.45 [0.31, 0.58]). Annual trophic position among female brown bears was therefore highly repeatable over multiple years ($R_{female}$ = 0.45 [0.31, 0.58]), indicating high between individual and low within individual variation in trophic position in females. This was corroborated by a strong correlation between the observed trophic position in a given year and the modelled posterior mean trophic position (Pearson correlation coefficient $r = 0.78$, $t = 22.63$, $df = 336$, $p < 0.001$, **Fig S3B**). In contrast, trophic position was only moderately repeatable in male brown bears ($R_{male}$ = 0.34 [0.18, 0.49], **Fig S3C**). After controlling for age, average trophic position estimated as the mean of the posterior distribution of the trophic position for each random intercept level ranged from 2.64 to 3.05 for individual females and from 2.9 to 3.38 for individual males (**Fig S3C**).

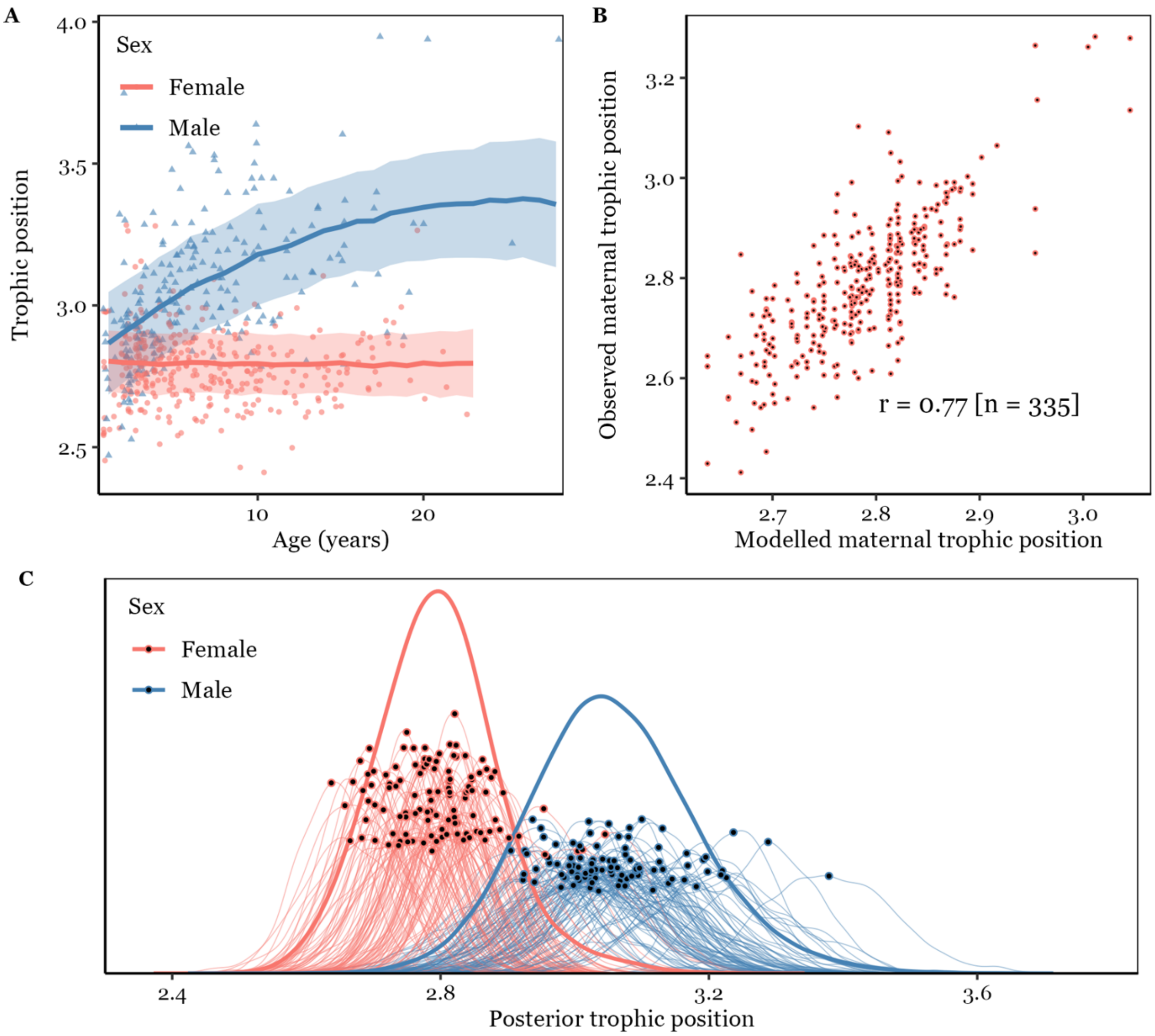


**Figure S3. A**) Effect of age (years) on trophic position in female (red) and male (blue) Scandinavian brown bears. Male bears increased in trophic position with age. For female bears there was no relationship between trophic position and age. **B)** We compared the observed maternal trophic position with the modelled maternal trophic position (i.e. the median of the posterior distribution). Based on 335 observations, observed and modelled trophic position were strongly correlated (r = 0.77). **C**) Posterior distribution of male and female, trophic position and each individuals' trophic position. The mean of the posterior distribution for each individual is indicated with black dots and was taken forward as maternal or paternal trophic position in the analysis presented in the main text.

**Table S1.** Posterior means with lower and upper bounds of 95% credible intervals of fixed effects in a linear mixed effects model evaluating the effects of age on annual trophic position in 104 female and 92 male brown bears.

| Covariate | mean | lowerCI | upperCI |
|---|---|---|---|
| Males (n = 219) | | | |
| Intercept | 3.06 | 3.02 | 3.10 |
| Age | 1.85 | 1.34 | 2.30 |
| Age.quadratic | -0.43 | -0.89 | -0.03 |
| Females (n = 335) | | | |
| Intercept | 2.80 | 2.78 | 2.82 |
| Age | -0.04 | -0.32 | 0.29 |
| Age.quadratic | 0.03 | -0.23 | 0.29 |

### 4. Maternal learning during rearing

We assume that daughters may in part learn dietary specialization from their mother because they constantly forage together during their first year of life. For this we look at the effect of maternal trophic position on daughter trophic position after independence. We here validated that maternal and daughter trophic position were correlated during rearing, providing the basis for a maternal learning effect after independence. We obtained records of maternal trophic position during rearing and daughter trophic position at age 0 (at the time of rearing) for 116 mother-offspring pairs (**Fig S4**). The correlation between the observed maternal trophic position and the observed daughter's trophic positions was strongly positive (Pearson correlation coefficient $r = 0.66$, $t = 9.36$, $df = 114$, $p < 0.001$).

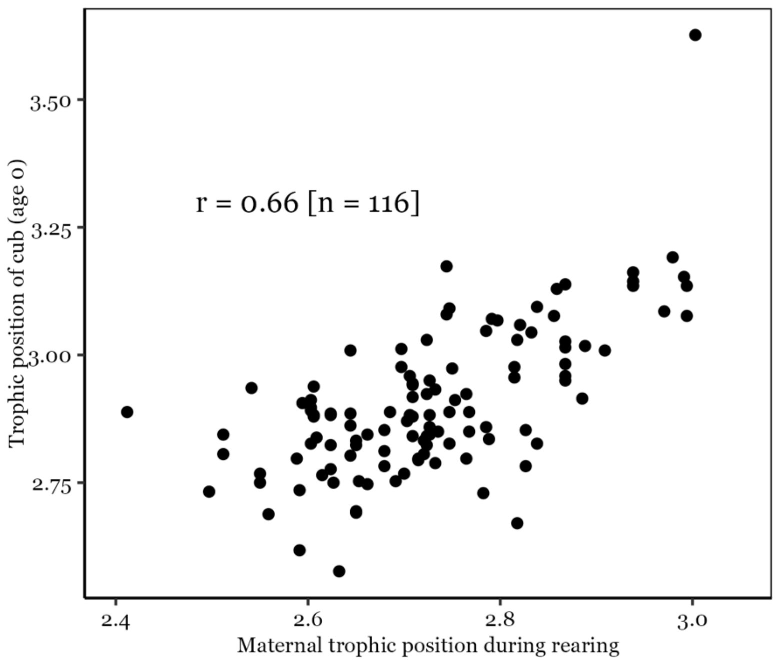


**Figure S4.** Relationship between maternal and daughter trophic position during rearing (i.e. the offspring's first year of life).

### 5. Distance between mother and daughter home range centroids over time and changes in home range scale environmental composition through time

Individuals living close in space may share a similar environment with access to similar resources and therefore display a similar dietary specialization. Indeed, both environmental similarity, i.e., "likeness" of home range habitat composition (main text) and spatial proximity (**Supplement 6**) explain similarity in dietary specialization. We found that the maternal trophic position was strongly correlated with the offspring trophic position shortly after family breakup but this correlation faded over time. We interpreted this effect as social maternal learning which fades over time, however, if daughters disperse over time (see **Figure S5A**) a fading social maternal learning effect could be confounded with an increasing spatial distance between mothers and daughters and co-occurring changes in resource availability.

We here explore how the distance between the natal range and the offspring's home range centroid changes over the years after separation (**Fig S5A**). Home range centroids were calculated on the basis of GPS or VHF locations during the active season (see main text). Indeed, we found support that the distance between the daughter's and mother's home range centroid increased over time in a non-linear fashion. The distance increased in the first 3 years and then leveled off such that the estimated distance in the first year after family breakup was 3.4 km and in year six after breakup 6 km. Overall the median dispersal distance was 8 km (range 1.4 – 28.8 km). An increase in distance by 2.5km, as well as a dispersal distance of 8km is negligible in biological terms of habitat composition or resource abundance given that moose are a highly mobile species. In addition, in **Supplement 6** we show that spatial autocorrelation in habitat composition seizes after 10 - 15km.

Alternatively, resource abundance within home ranges could also change over the lifetime of individuals if habitat composition changes. Though our Corine Landcover Map was static, we accounted for forestry activity by manually updating the map with newly emerging clearcuts, based on annual data of forest clearcutting by the Swedish Forestry Agency (Skogsstyrelsen). We manually recategorized mature forest stands to clearcuts in years when they were harvested; 9 years after harvest we recategorized them as young forests and 20 years after harvest as mature stands. As environmental similarity is fitted with as constant environmental similarity matrix, we here provide a validation that the proportion of forest cover (**FigS5B**) and disturbed forest (**FigS5C**) in a bear's home range only shifted minorly over the lifetime of an individual. We cannot exclude the possibility that we missed to account for unregistered changes in habitat composition, in particular forest disturbance.

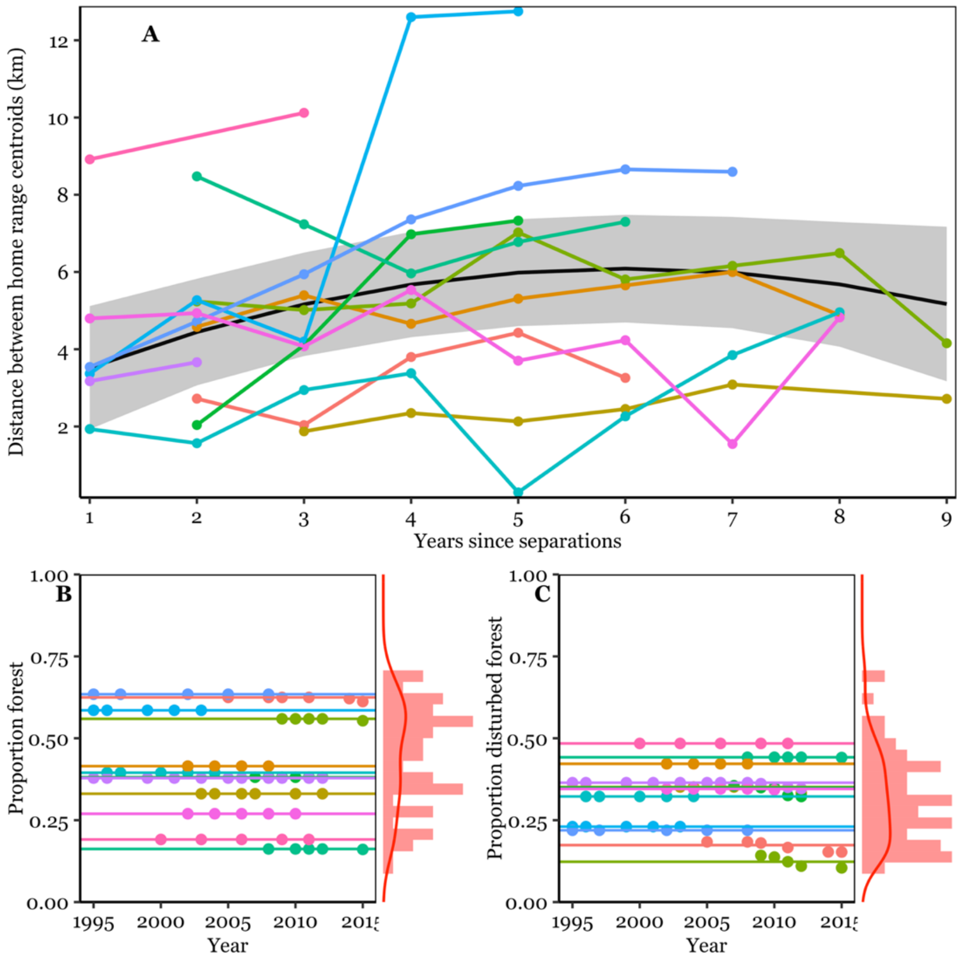


**Figure S5. A)** Distance between the natal range centroid and the daughters home range centroid after family breakup for 12 female brown bears (i.e., daughters) monitored for up to 9 years after separation. The distance between the natal range centroid and the daughter's home range centroid increases by about 2.5 km in the first years after separation but keeps stable later on. Home range composition changed little over the lifetime of individuals (**B & C)** corroborating that constant estimates of environmental similarity used in our analysis are robust.

### 6. Effect of spatial distance on trophic position

In the main text of the manuscript, we tested whether variation in environmental habitat composition drives individual variation in trophic position by accounting for pairwise environmental similarity (proportion of forest, proportion of disturbed forest and habitat diversity). We used correlograms to assess the spatial autocorrelation of the proportion of habitat metrics within lifetime home ranges at different pairwise spatial distances (using home range centroids). Spatial autocorrelation seized after 10-15km, which was farther than the median dispersal distance of 8km (**Fig S6 A**). While environmental similarity incorporates aspects of habitat composition in the environment, it does not account for unmapped aspects such as resource density. Where resources change in a continuous fashion across the study area for reasons other than habitat composition, environmental similarity may not accurately reflect resources availability. For example, ant or berry availability will be tightly linked to the availability of suitable habitats, while moose density (a highly mobile species) will likely change over larger spatial scales and could be better captured using pairwise spatial distance among bear home ranges rather than environmental similarity. On the other hand, female brown bears are highly philopatric potentially leading to confounds between spatial distance and maternal effects. Therefore, bears inhabiting home ranges close in space may display a more similar trophic position, either because they experience more similar resource availabilities or because female philopatry leads to clusters of mothers and daughters in space (so called matrilines) which may display similar trophic positions due to social learning or shared genes.

We here present an alternative analysis where we substitute the environmental similarity matrix with a spatial distance matrix (S matrix) (Gervais *et al.* 2022). We first calculated lifetime individual home ranges using the 95% kernel density estimator including relocation data of all years a bear was monitored post separation from the mother. We included bears with at least 1000 GPS locations or VHF locations obtained on at least 25 days to construct home ranges and from these, extracted lifetime home range centroids. We calculated a pairwise Euclidian distance matrix between all bear home range centroids to account for spatial autocorrelation driven by spatial proximity of home ranges. We then refit the *fixed* model from the main text including a second order fixed effect for age (fitted as the main effect of time since separation of the mother), and maternal trophic position interacting with time since separation and partitioning the remaining variance into spatial distance ($V_S$), additive genetic ($V_A$), maternal ($V_M$), permanent between-individual ($V_I$), and residual within-individual effects ($V_R$).

Although spatial distance and environmental similarity were significantly correlated, the correlation coefficient was too weak to indicate relevant spatial autocorrelation (Mantel test: Spearman's rho = 0.109, p = 0.003). Environmental similarity accounted for 9% (0.1 – 49%) of the variance in trophic position while spatial proximity accounted for 50% (3 – 80%) of the variance in trophic position (**Fig S6B**). Bears living in close proximity therefore have more similar diets than bears inhabiting home ranges farther apart. Habitat composition does not explain this spatial variation in diet well suggesting that either resource availability varies across the landscape for factors other than habitat composition or that diet is spatially autocorrelated for other reasons than just resource availability. For example, due to philopatry, closely related individuals inhabit home ranges closer in space. Indeed, maternal effects and social maternal learning explain more variance in the environmental similarity model than in the spatial distance model suggesting that spatial distance is indeed correlated with female social organization.

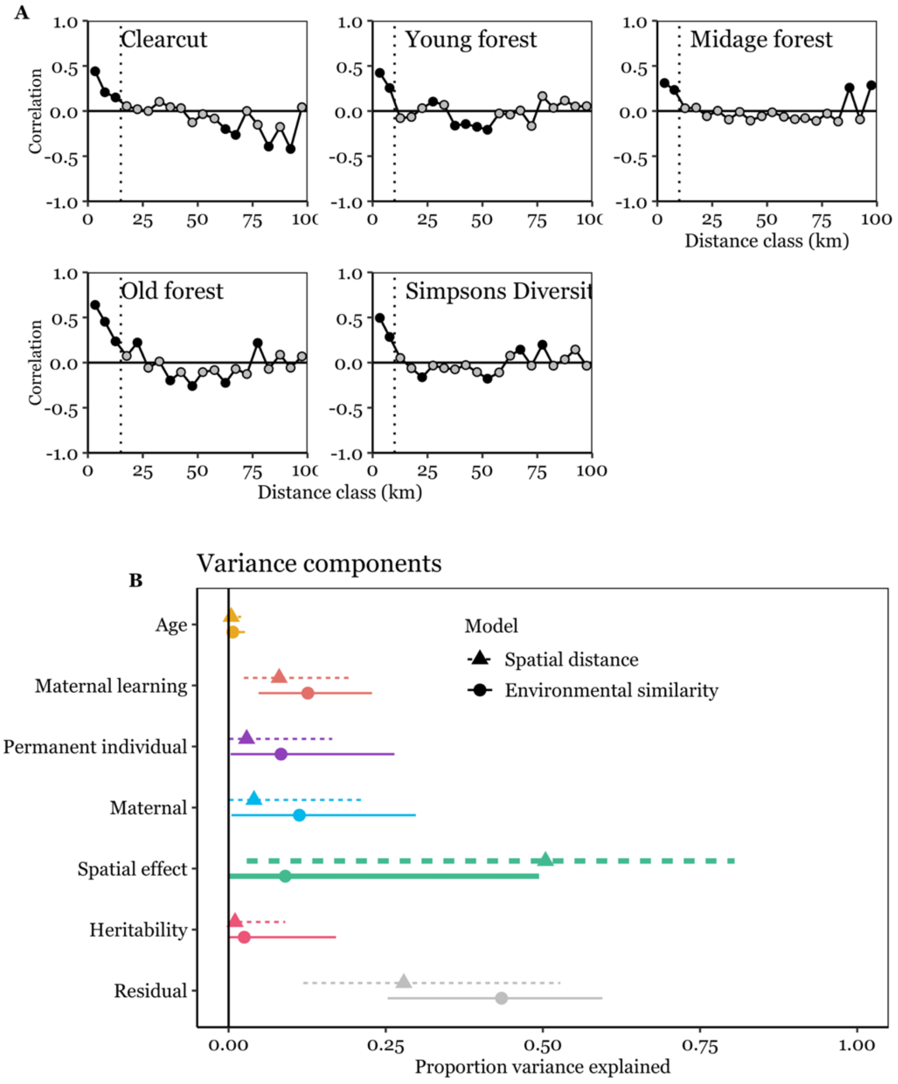


**Figure S6. Figure S6. A)** Correlograms showing spatial autocorrelation for habitat metrics measured as the proportion of cover in bear home ranges seizes after 10-15 km, **E)** Proportion of variance explained (median and 89% equal tails intervals) by a model controlling for spatial effects on variation in bear diet with pairwise spatial proximity among bear home range centroids (dotted line and triangle) or environmental similarity (solid line and filled dot, also see main text). Environmental similarity was defined as the percentage of old and mid-aged forest habitat, disturbed forest habitat, and habitat diversity (Simpson's diversity index) in a bear's home range (95% kernel utilization distribution). Proximity of home ranges explained more variance than similarity in habitat composition suggesting that other factors than habitat per se drive spatial variation in resource abundance.

### 7. Collinearity of genetic and permanent maternal effects with spatial proximity

We here assess whether genetic effects on trophic position (where more closely related individuals also have a more similar trophic position) and/or maternal effects (where siblings of the same mother have a more similar trophic position) are confounded by closely related individuals also living in close proximity to each other and hence sharing the same environment. To this end, we compared the model controlling for environmental effects with a *spatial distance* matrix (**supplementary analysis 6, Fig S6**) with a reduced model without the spatial distance covariance matrix (*no spatial distance model*) and with a further reduced model also excluding the maternal effect (*no spatial distance + no maternal model*).

In both reduced models, no great increase in the variance explained by genetic effects (i.e. heritability) could be detected (narrow-sense heritability *spatial distance model* = 1% [0.01 – 9%]; *no spatial distance model* = 4% [0.05 – 20%]; *no spatial distance + no maternal model* = 5% [0.05 – 27%]; **Fig S7** pink lines). Instead, the variance was mostly attributed to maternal learning (increase from a median 8% to a median of 15% in both reduced models, respectively), maternal effects (increase from a median 4% to a median of 13% in the *no spatial distance model*), and permanent individual effects (increase from a median of 3% to a median of 11% and 22%, resp.). Our results suggest that maternal effects (both maternal learning and maternal effects) but not heritability *per se* are partially masked by spatial distance, because daughters settle close to their maternal home range forming clusters of closely related females over time (matrilines). We are therefore confident that results presented in the main manuscript, where we control for habitat composition and not spatial distance, provide less confounded estimates allowing to attributed variance in dietary specialization to its biological sources.

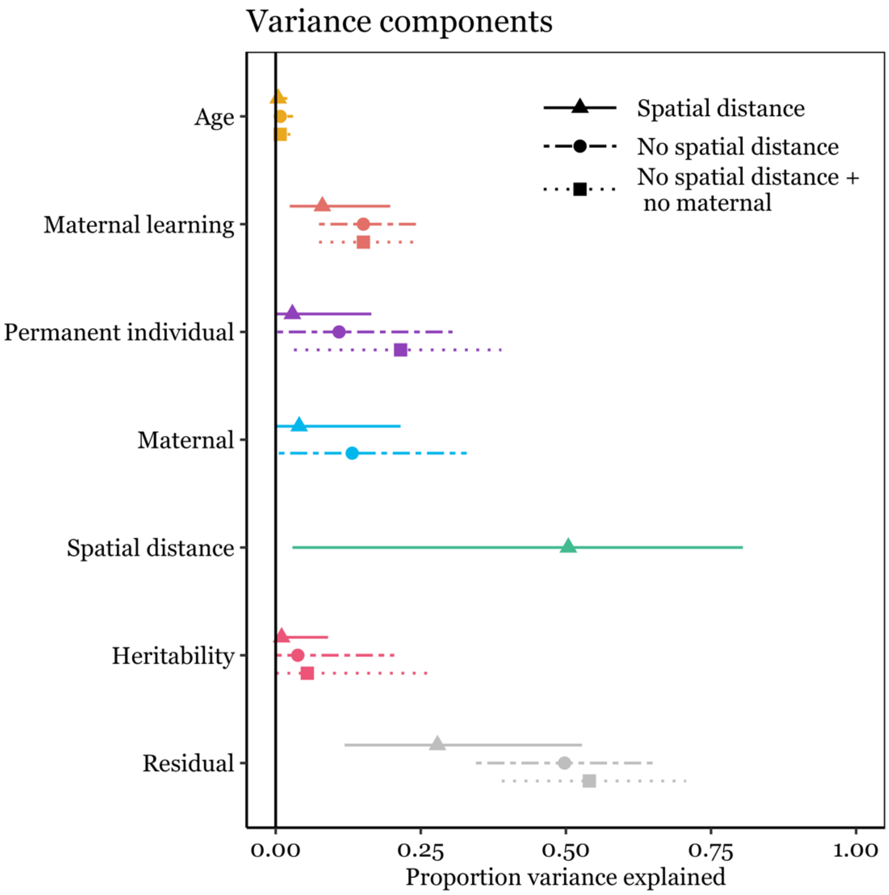


**Figure S7.** Proportion of variance explained in the *spatial distance* model (triangle and dotted line) by permanent among-individual, spatial proximity, additive genetic, permanent maternal, and residual within-individual effects. Narrow-sense heritability did not increase significantly in a model excluding the potentially confounding effect of spatial proximity (dot and solid line), instead variance attributed to maternal learning, maternal effects, and permanent individual effects increased.

### 8. Effect of maternal trophic position on male and female offspring

The analysis in the main text identified significant time related changes in the correlation between maternal and offspring trophic positions. In particular, the effect of maternal trophic position on daughter trophic position seized approximately 5 years after family breakup (see Figure 3 in the main text). The offspring sex ratio in our population is even but male offspring are not monitored throughout life because they usually disperse out of the study area. Because social maternal learning which affects diet composition in the first 4 years after separation may not be limited to female offspring, we evaluated whether this relationship also holds true in male offspring. We identified 31 male and 69 female offspring with known maternal trophic position (n = 36 unique mothers). Using offspring trophic positions, measured at the earliest time within the first 4 years after separation as response variable (no repeated measures per individual) we fit a simple linear regression, evaluating the effects of maternal trophic position, sex of the offspring (male or female) and their interaction. Neither the interaction term (AIC = -89.87), nor the main term of sex (AIC = -91.73) improved the model and the simple model with only maternal trophic position as predictor ($\beta_{MaternalTP} = 1.12 \pm 0.23$) was selected as the most parsimonious model (AIC = -93.03). Maternal trophic position alone explained 19% of the variance in offspring trophic position in the first four years of independence. The Pearson correlation coefficient was 0.45 ($t = 4.12$, $df = 67$, $p < 0.001$) between maternal and female offspring trophic position (n = 69 daughters) and 0.4 ($t = 2.36$, $df = 29$, $p = 0.03$) between maternal and male offspring trophic position (n = 31 sons) in the first 4 years after family breakup.

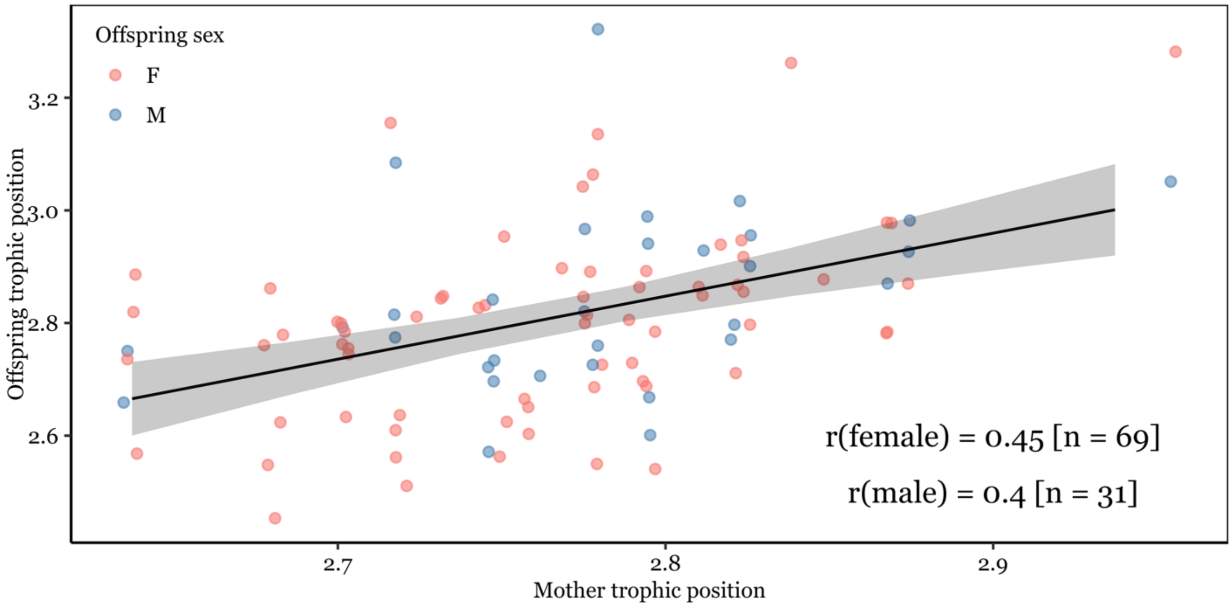


**Figure S8.** Offspring trophic position was affected by maternal trophic position but was unaffected by offspring sex in the first 4 years after family breakup. Maternal trophic position explained 19% of the phenotypic variance in offspring trophic position. The Pearson correlation coefficient between maternal trophic position offspring trophic position was 0.45 for females and 0.4 for males.

#### 9. Effect of paternal trophic position on offspring trophic position

For 49 individuals sired by 25 unique fathers, we also obtained information about the paternal trophic position, allowing us to draw some preliminary conclusions about the importance of paternal effects. Choosing the earliest measure per individual after separation from the mother we found no relationship between the fathers and offspring trophic position (Pearson correlation coefficient r = 0.13, t = 0.87, df = 47, p = 0.39).

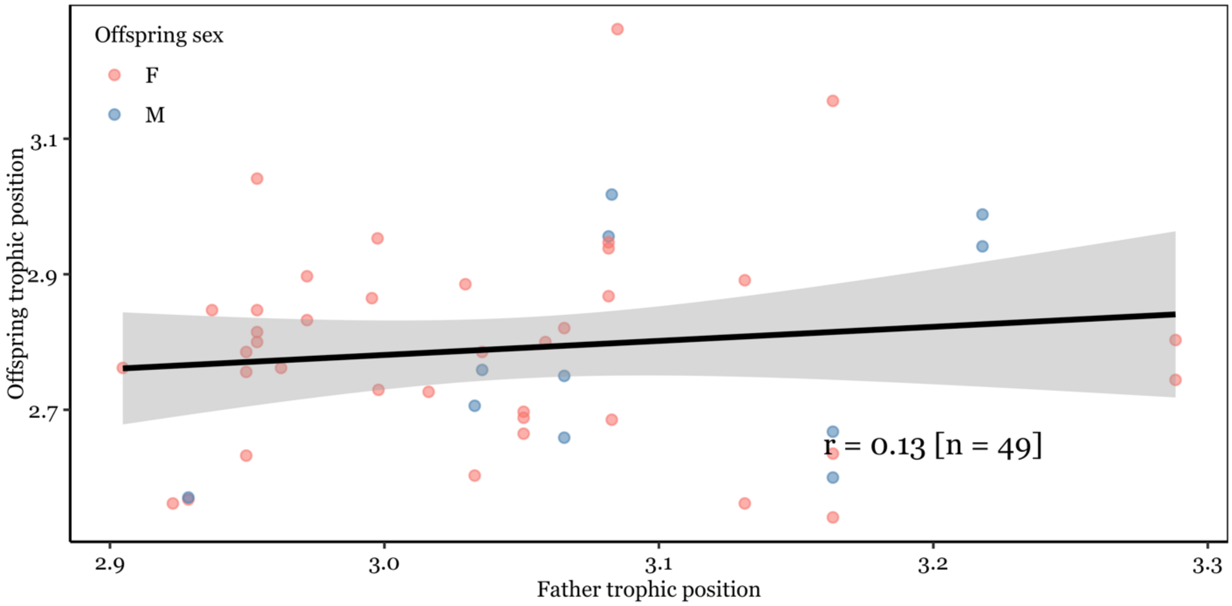


**Figure S9.** Single measurements of offspring trophic position (n = 46) were not correlated with the trophic position of the father (n = 23 unique fathers).

#### 10. Alternative analysis of the effect of maternal learning

The analysis in the main text identified a significant effect of maternal learning, defined as a phenotypic correlation of a daughters and a mother trophic position after separation. We modelled maternal trophic position as the mean of the posterior distribution of a mother's random intercept in a population wide model (**S3**), where 45% of the total phenotypic variance in trophic position could be attributed to permanent individual effects and were generally strongly correlated (r = 0.78, **FigS3B**). Yet, arguably, using the observed trophic position of the mother during rearing (i.e. in the first year of an offspring's life) would provide the most accurate measure of any maternal effect. We here replicate the main analysis by included the observed maternal trophic position during rearing instead of the modelled trophic position to a reduced dataset of 62 records from 38 individuals (instead of 213 records from 71 individuals). The reduction in sample size precludes us from fitting a complex model as shown in the main manuscript and we only fit the fixed effects structure of maternal trophic position interacting with a nonlinear effect (second order polynomial) of time since separation and controlling permanent individual effects with a random intercept for BearID.

The alternative model corroborated the absence of an age effect (explained variance = 1.8% [0% - 7%]) and strong permanent individual variation (explained variance = 46% [6% - 73%]) in the same magnitude as found in the main text and S3. At the same time, maternal learning had an even stronger effect (22% [8% - 37%]) in shaping dietary specialization than found in the main analysis (13% [5% - 23%]). We therefore conclude that using modelled posterior maternal trophic positions provide an adequate measure for estimating the effect of maternal diet on her female offspring's diet later in life. However, using modeled maternal positions seems to provide a conservative estimate of maternal learning and may in fact underestimate the true effect of maternal learning.

### 11. Summary table of full model

**Table S2.** Summary table of the final model presented in the manuscript: Posterior medians with lower and upper bounds of 89% equal tails credible intervals of fixed effects and variance components in a linear mixed effects model evaluating drivers of dietary specialization, measured as trophic position in female brown bears. Estimates of variance are given in standard deviations.

| | Median | Upper 89% ETI | Lower 89% ETI |
|---|---|---|---|
| Fixed effects | | | |
| Intercept | 2.813 | 2.733 | 2.909 |
| Maternal trophic position (TP) | 0.042 | 0.014 | 0.071 |
| Time separated (1) | 0.12 | -0.093 | 0.348 |
| Time separated (2) | -0.101 | -0.305 | 0.096 |
| maternalTP : Time separated (1) | -0.534 | -0.715 | -0.335 |
| maternalTP : Time separated (2) | 0.125 | -0.019 | 0.271 |
| Variance components | | | |

| | | | |
|---|---|---|---|
| $sd_{PermanentIndividual}$ | 0.047 | 0.01 | 0.085 |
| $sd_{AdditiveGenetic}$ | 0.027 | 0.003 | 0.067 |
| $sd_{EnvironmentalSimilarity}$ | 0.048 | 0.005 | 0.144 |
| $sd_{MaternalEffects}$ | 0.055 | 0.011 | 0.096 |
| residual | 0.107 | 0.095 | 0.119 |

The complete model output with raw posterior distributions can be accessed in the data & code repository: https://doi.org/10.17605/OSF.IO/68B9U.

### 12. Statistical power

Estimating heritability from additive genetic variance requires larger sample sizes than simple mixed models because additive genetic variance in estimated from a covariance matrix including all pairwise comparisons of individuals and not simple grouping factors (i.e., random intercept). A recent analysis, estimating additive genetic variance and heritability of fitness across 19 populations in the wild included datasets with a minimum of 880 individuals (Bonnet *et al.* 2022). Our study has a pedigree spanning 1640 individuals but our study relied on field observations of mother-daughter pairs and our final sample size was 72 individuals, we therefore likely have low statistical power to estimate additive genetic variance. Indeed, asymmetric posterior distributions of variance components, which is the case for our model (**Fig S6E**), are indicative of low statistical power.

Power analysis is a concept rooted in frequentist statistics and estimates the probability to reject the null hypothesis given the sample size (i.e., to detect that an effect is statistically significantly different from 0 at a prescribed alpha level, usually 0.05). Estimating power for variance components is ambiguous because variances are bound by 0. Elsewhere, significance of variance components has been determined using its adaptation potential over a few generations, where attributed variances < 0.001 were considered small (Bonnet *et al.* 2022). Pick *et al.* (2023) recently suggested an alternative approach where significance can be estimated by comparing the variance explained by a given component to a null distribution. This null distribution can be either simulated or achieved through permutation of the original dataset.

We here used the permutation approach to generate a p-value for additive genetic variance. We kept the architecture of our pedigree (Dam/Sire pairs) but randomly assigned parents to offspring. If related individuals have similar dietary specialization (i.e., heritability of dietary specialization), our observed pedigree should explain more variance than a permutated pedigree. We generated a p-value by calculating the proportion of permutated models where the explained variance is larger than the explained variance in the observed dataset. We fitted a reduced "basic animal model", controlling only for fixed effects, bear ID and genetic structure. We permutated our dataset 1000 times, fitting 1000 animal models as null distribution. Our observed genetic pedigree did not explain more variance than the permutated null distribution ($p = 0.077$).
